# Microfluidic Co-Culture for Modeling Human Joint Inflammation in Osteoarthritis Research

**DOI:** 10.1101/2025.01.27.635089

**Authors:** Hosein Mirazi, Scott T. Wood

## Abstract

Here we present a microfluidic model that allows for co-culture of human osteoblasts, chondrocytes, fibroblasts, and macrophages of both quiescent (M0) and pro-inflammatory (M1) phenotypes, maintaining initial viability of each cell type at 24 h of co-culture. We established healthy (M0-based) and diseased (M1-based) joint models within this system. An established disease model based on supplementation of IFN-γ and LPS in cell culture media was used to induce an M1 phenotype in macrophages to recapitulate inflammatory conditions found in OA. Cell viability was assessed using NucBlue™ Live and NucGreen™ Dead fluorescent stains, with mean viability of 83.9% ± 14% and 83.3% ± 12% for healthy and diseased models, respectively, compared with 93.3% ± 4% for cell in standard monoculture conditions. Cytotoxicity was assessed via a lactate dehydrogenase (LDH) assay and showed no measurable increase in LDH release into the culture medium under co-culture conditions, indicating that neither model promotes a loss of cell membrane integrity due to cytotoxic effects. Cellular metabolic activity was assessed using a PrestoBlue™ assay and indicated increased cellular metabolic activity in co-culture, with levels 5.9 ± 3.2 times mean monolayer cell metabolic activity levels in the healthy joint model and 5.3 ± 3.4 times mean monolayer levels in the diseased model. Overall, these findings indicate that the multi-tissue nature of *in vivo* human joint conditions can be recapitulated by our microfluidic co-culture system at 24 h and thus this model serves as a promising tool for studying the pathophysiology of rheumatic diseases and testing potential therapeutics.

## Introduction

Osteoarthritis (OA) is a crippling health condition generally characterized by chronic joint pain, degeneration of articular cartilage, synovitis, and bone remodeling. (Tong et al., 2022) OA affects over 520 million people worldwide, and frequently progresses to the point of requiring joint replacement in late stages of the disease because no disease-modifying OA drugs have yet received regulatory approval. (Grässel & Muschter, 2020; Long et al., 2022) An important limitation of most prior studies is that, as illustrated in Figure 1, the monoculture models used in those studies are devoid of cellular crosstalk, which is an important constituent of the joint as an organ. (Loeser et al., 2012) The development of preclinical *in vitro* models reproducing the complexity of human joint diseases constitute an important trend in OA research, yet current models are still limited in their capacity to recapitulate the complex multicellular nature of human joints, hindering the development of an efficient therapy for OA. (Cope et al., 2019; Domínguez-Oliva et al., 2023; Malfait & Little, 2015; Perisin & Sund, 2018; Swearengen, 2018; Xie et al., 2018)

**Figure 1:**
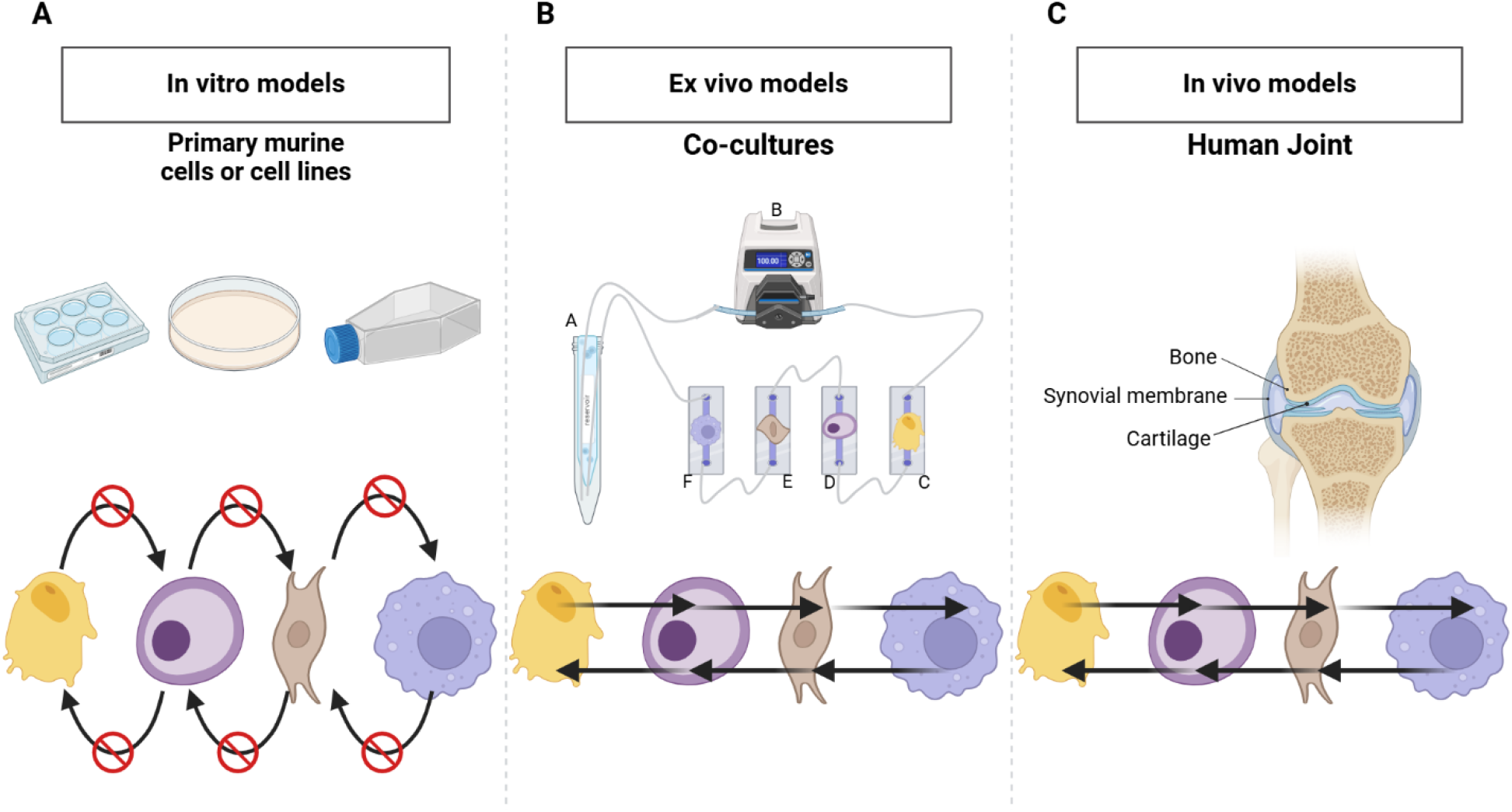
Comparison of Monoculture, Co-culture and In Vivo Human Joint Systems. (**A)** classic monoculture model in static conditions-does not provide the possibilities for crosstalk among the cell types present, which limits cellular interaction. **(B)**, Co-culture model developed during this work that consists of more than one cell type in communication via shared cell culture media hence active crosstalk/interactions. **(C)** Human joint in vitro model with bone-synovial membrane-cartilage, whose tissues normally interact and signal with each other for the maintenance of function and responsiveness. Created in BioRender. Mirazi, H. (2025) https://BioRender.com/j17f652.

*In vivo* models, though providing valuable insights into OA, often fail to accurately mimic human physiology due to limitations in replicating the complex interactions between joint tissues. (Cope et al., 2019) This often leads to inconsistent and non-predictive outcomes in drug development studies. (Dou et al., 2023; Guo et al., 2022; Zaki et al., 2022) Existing *in vitro* models, while more advanced, are generally restricted to the cartilage compartment, and co-culturing of key cell types involved in OA—such as osteoblasts, chondrocytes, fibroblasts, and macrophages—is uncommon. (Banh et al., 2022; Haltmayer et al., 2019; He et al., 2020; Pirosa et al., 2021; Salgado et al., 2021) One major challenge contributing to this gap is that these different cell types have distinct media requirements, making it difficult to maintain all cells under the same conditions. (Weiskirchen et al., 2023) Additionally, some multi-cell models rely on induced pluripotent stem cells (iPSCs) instead of using patient-derived cell types representative of mature cells from each specific tissue, which limits the accuracy of disease representation. (Li et al., 2023; Makarcyzk et al., 2023)

Unlike classical mono-culture models, co-culture systems allow interactions to occur between cell types, a situation much more similar to the *in vivo* environment of human joints. The use of advanced bioengineered models, such as microphysiological systems (MPSs), which includes organoids and organs on chips, likely represents the future of OA research. (Banh et al., 2022; Goers et al., 2014; Hofer & Lutolf, 2021) These systems are designed to capture these intercellular interactions and provide a more physiologically relevant translation of human physiology than earlier approaches. (McNerney et al., 2021; Palasantzas et al., 2023; M. Rothbauer et al., 2021) Figure 2 illustrates different approaches to disease modeling, including the use of immune cells, such as macrophages, to simulate inflammatory conditions. A model with the involvement of all the key cell types of a human joint-osteoblasts, chondrocytes, fibroblasts, and macrophages-could mark a pivotal advance leading to fundamental changes in our understanding of the pathogenesis of OA and other complex joint diseases.

**Figure 2:**
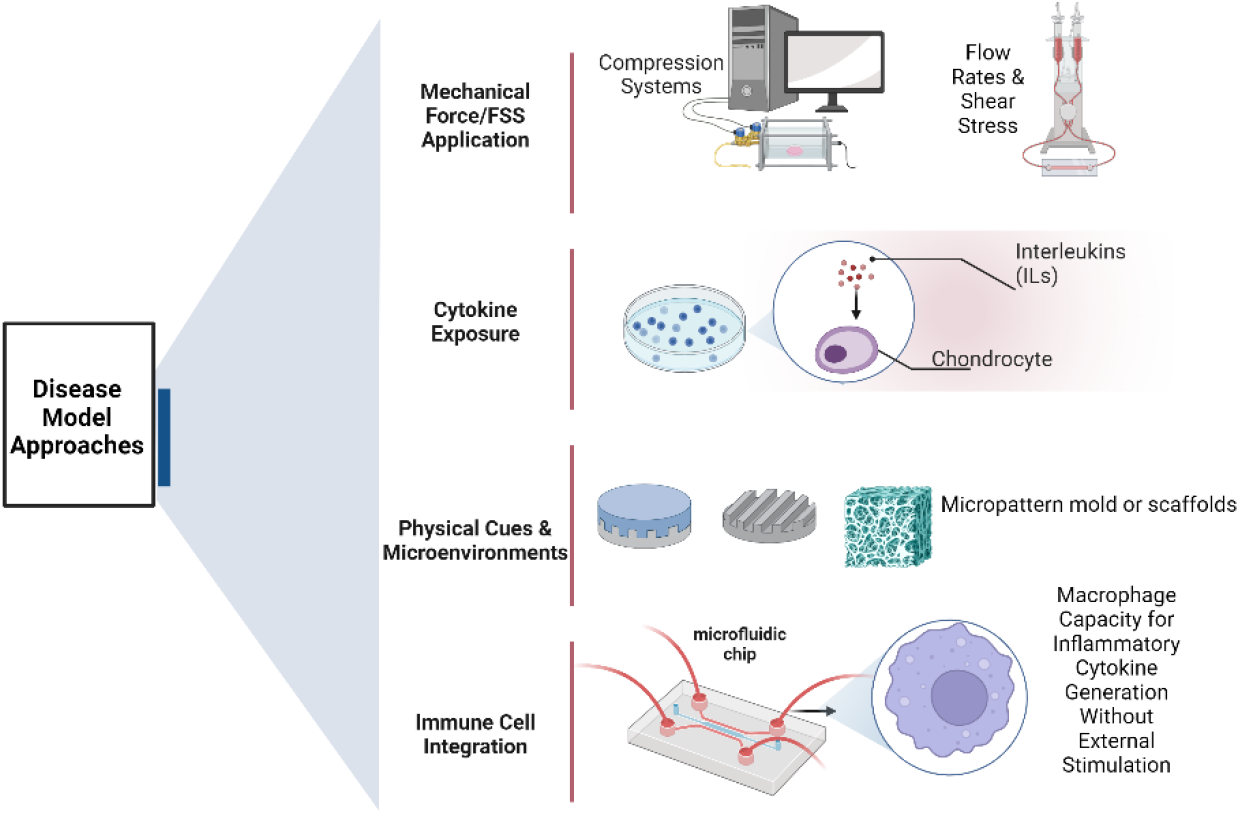
Methods of disease model development. **(A)** group of methods relies on flow rates or applies mechanical forces/fluid shear stresses as a method of stimulation to the cells. **(B)** Inflammatory cytokines, such as interleukins or other stimulators, are used directly on the cells for inducing disease states. **(C)** Physical stimuli or microenvironments-for example, various scaffolds-are used with the express intention of directing and eliciting from the cells responses that are characteristic of diseases. **(D)** To simulate conditions seen in inflammatory diseases, the system is treated with immune cells, including macrophages, which stimulate the secretion of inflammatory cytokines. Created in https://BioRender.com

The most critical limitation of most of the multi-cell models available, especially the ones based on iPSCs, is that they generally either do not differentiate precisely into the highly specialized cell types that are critical for studying the osteoarthritic joint environment, most notably adult articular chondrocytes. As most iPSC-based models are cytokine-driven, their differentiation is susceptible to be driven to inhomogeneous and non-physiological phenotypes, and hence it can be difficult to accurately recapitulate key characteristics of cells participating in OA pathogenesis. (Kao et al., 2023; Zhou et al., 2021) Some co-culture models have been published that include a limited number of human joint cell types, but to date, no studies have been published establishing co-culture conditions to mimic the joint microenvironment using cells from three different joint tissues, the most universal of which are bone, cartilage, and synovium. (Awad et al., 2023; Kao et al., 2023) This has created a need for an integrated system bringing together cells from these essential components of the joint for capturing the multifactorial nature of OA, and of the joint as an organ, more effectively.

To address the challenges associated with co-culturing bone, cartilage, and synovium cells together, we utilized a microfluidic co-culture system designed to replicate the complex paracrine dynamics of the human joint environment. Unlike traditional static models, our system enables these distinct cell types to interact under controlled flow conditions using a combined cell culture media, allowing us to overcome the limitations of differing media requirements. The goal of this study was to create a biologically relevant platform that integrates osteoblasts, chondrocytes, fibroblasts, and both quiescent (M0) and pro-inflammatory (M1) macrophages as illustrated in Figure 3, thereby mimicking paracrine cellular interactions under both healthy and diseased conditions. We hypothesized that, on average, cell viability and metabolic activity could be maintained at ≥70-80% of baseline monoculture levels through this shared microenvironment that promotes paracrine crosstalk between cell types. By successfully replicating these paracrine interactions using human patient-derived cells, we provide the foundation for a novel system capable of advancing the understanding of OA pathogenesis and enabling the testing of therapeutic interventions in a reliable and replicable manner.

**Figure 3:**
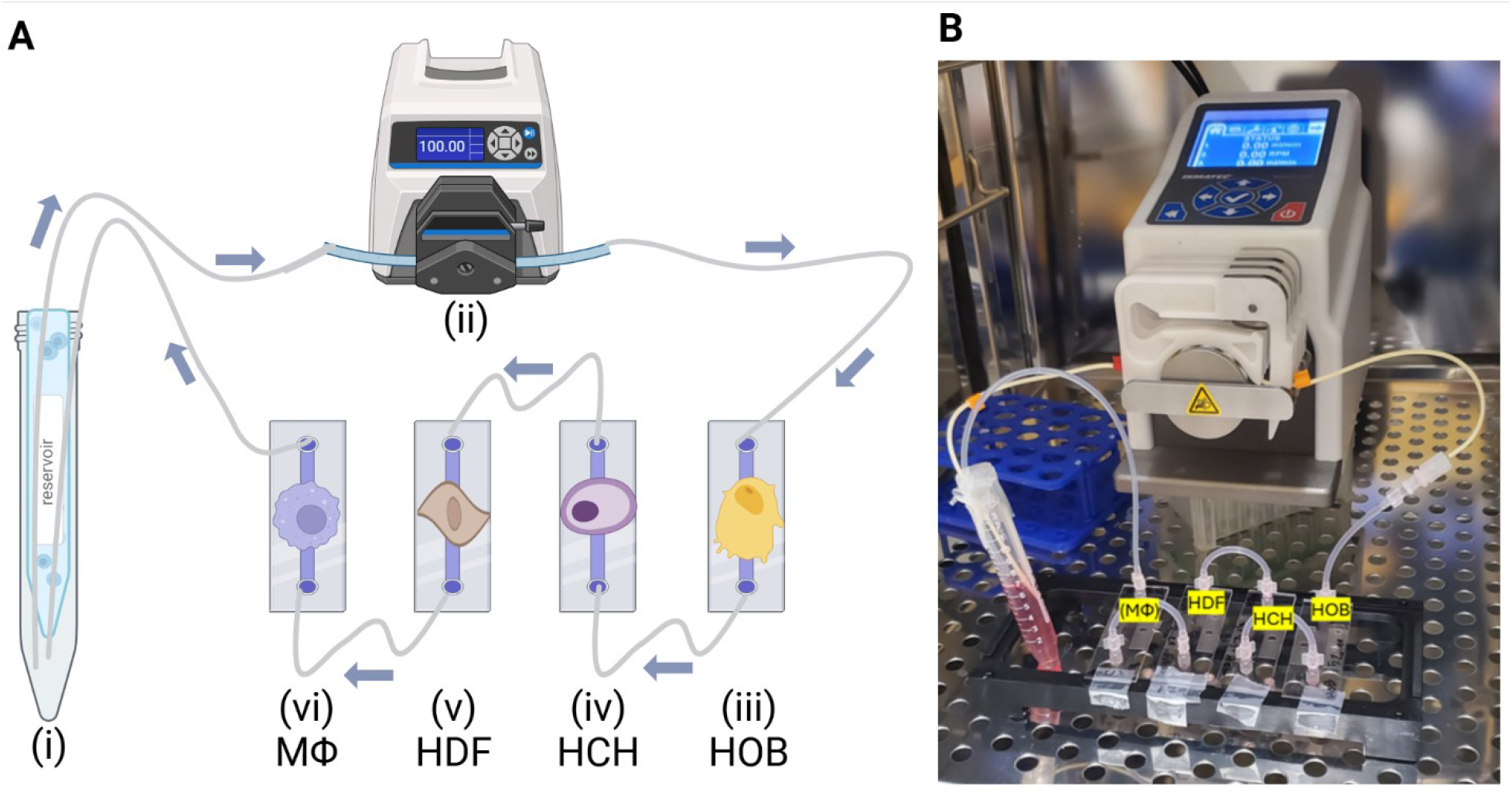
Experimental Setup and Workflow. **(A)** Illustration and **(B)** Photo of experimental setup placed inside the incubator, where the microfluidic system operated under controlled conditions for the duration of the experiment. Created in BioRender. Mirazi, H. (2025) https://BioRender.com/u04h894

## Materials and Methods

### Cell Culture

Primary human osteoblasts (HOBs) isolated from the cancellous bone of the femoral head of an 85-year-old Caucasian female were obtained from PromoCell (Heidelberg, Germany) and cultured per manufacturer recommendations. Briefly, HOBs were expanded in Osteoblast Growth Medium in T-75 flasks at 37°C in a 5% CO2 humidified incubator and used between passages four to six. Following expansion, HOBs were seeded in microchannels at a density of 40,000 cells/cm2 for experimentation.

Primary human chondrocytes (HCHs) isolated from the cartilage of the tibial head of a 74-year-old female were obtained from PromoCell and cultured per manufacturer recommendations. HCHs were expanded in Chondrocyte Growth Medium in T-75 flasks at 37°C in a 5% CO2 humidified incubator and used between passages three to five. Following expansion, HCHs were seeded in microchannels at a density of 80,000 cells/cm2 for experimentation.

Primary human dermal fibroblasts (HDFs) isolated from adult skin (female, 49 years old) were obtained from the American Type Culture Collection (ATCC) and cultured per manufacturer recommendations. Briefly, HDFs were expanded in Dulbecco’s Modified Eagle’s Medium (DMEM) supplemented with 10% fetal bovine serum (FBS) and 1% penicillin-streptomycin (P/S) in T-75 flasks, incubated at 37°C in a 5% CO2 humidified environment and used between passages four to six. Following expansion, HDFs were seeded in microchannels at a density of 40,000 cells/cm2 for subsequent experimentation.

THP-1 monocytes isolated from the peripheral blood of a one-year-old male patient with acute monocytic leukemia were obtained from ATCC and cultured per manufacturer recommendations. THP-1s were expanded in RPMI-1640 Medium supplemented with 10% fetal bovine serum (FBS), 1% penicillin-streptomycin (P/S), 2 mM L-glutamine, 10 mM (2-[4-(2-hydroxyethyl) piperazin-1-yl] ethanesulfonic acid) (HEPES), 1 mM sodium pyruvate, 4.5 g/L glucose, and 1.5 g/L sodium bicarbonate. THP-1 cells were maintained in T-25 flasks at a seeding density of 4.0 × 105 viable cells/mL, reaching approximately 1.4 × 106 viable cells/mL after six days with media addition, under incubation at 37°C in a 5% CO2 humidified environment.

Prior to experimentation, THP-1 cells were differentiated into either the M0 (i.e., quiescent) or M1 (i.e., pro-inflammatory) macrophage (MΦ) phenotype, following the protocol previously described. (Baxter et al., 2020) Briefly, monocytes were seeded in microchannels at 75,000 cells/cm^2^, incubated with 50 nM Phorbol 12-Myristate 13-Acetate (PMA) for 48 hours, and then allowed to rest in PMA-free medium for an additional 24 hours to differentiate into THP-1-derived M0-type macrophages (M0MΦs). To induce the pro-inflammatory M1 phenotype, M0MΦs were treated for 24 hours with 20 ng/mL human interferon gamma (IFNγ) and 100 ng/mL lipopolysaccharide (LPS) derived from Escherichia coli O111.

### Microfluidic Co-Culture

A µ-Slide I Luer ibiTreat (Ibidi, Germany) system was utilized for cultivating osteoblasts, chondrocytes, fibroblasts, and macrophages under controlled flow conditions, regulated by a Masterflex® Ismatec® Reglo ICC Peristaltic Pump. Cells were seeded and allowed to attach overnight in a 5% CO2 incubator at 37°C. Once attachment was confirmed, the µ-Slide I Luer units were connected into a fluidic circuit, and shared media was circulated through the system for 2 hours, at a shear stress of 0.05 dyn/cm^2^ as illustrated in Figure 3. This process allowed thorough mixing, facilitating nutrient exchange and communication among the cells. After 2 hours of mixing under flow, circulation was stopped, and the cells were allowed to remain in the shared media for an additional 24 hours without active flow. This period aimed to enable paracrine interactions while also while also enabling media sampling specific to each cell type. Following the 24-hour static flow co-culture period, cell viability, cytotoxicity, and metabolic activity were assessed. These assays were selected to provide comprehensive insights into cell.

### Viability Assessment

Cell viability was assessed using a ReadyProbes™ Cell Viability Blue/Green Imaging Kit (Invitrogen). NucBlue™ Live stained all cell nuclei (displayed as green), while NucGreen™ Dead selectively labeled dead cells (displayed as magenta). When overlaid, live cells were visualized with green nuclei, dead cells appeared with white nuclei, and magenta-stained nuclei indicated problematic areas (Figure 4A). Samples were incubated with two drops of each reagent per mL of culture media for 20 minutes at 37°C. Fluorescence microscopy was performed using an Olympus IX71 microscope equipped with a UPLFLN 10X objective and an Andor iXon Ultra EMCCD camera (Andor) to capture high-resolution images for quantitative analysis.

**Figure 4:**
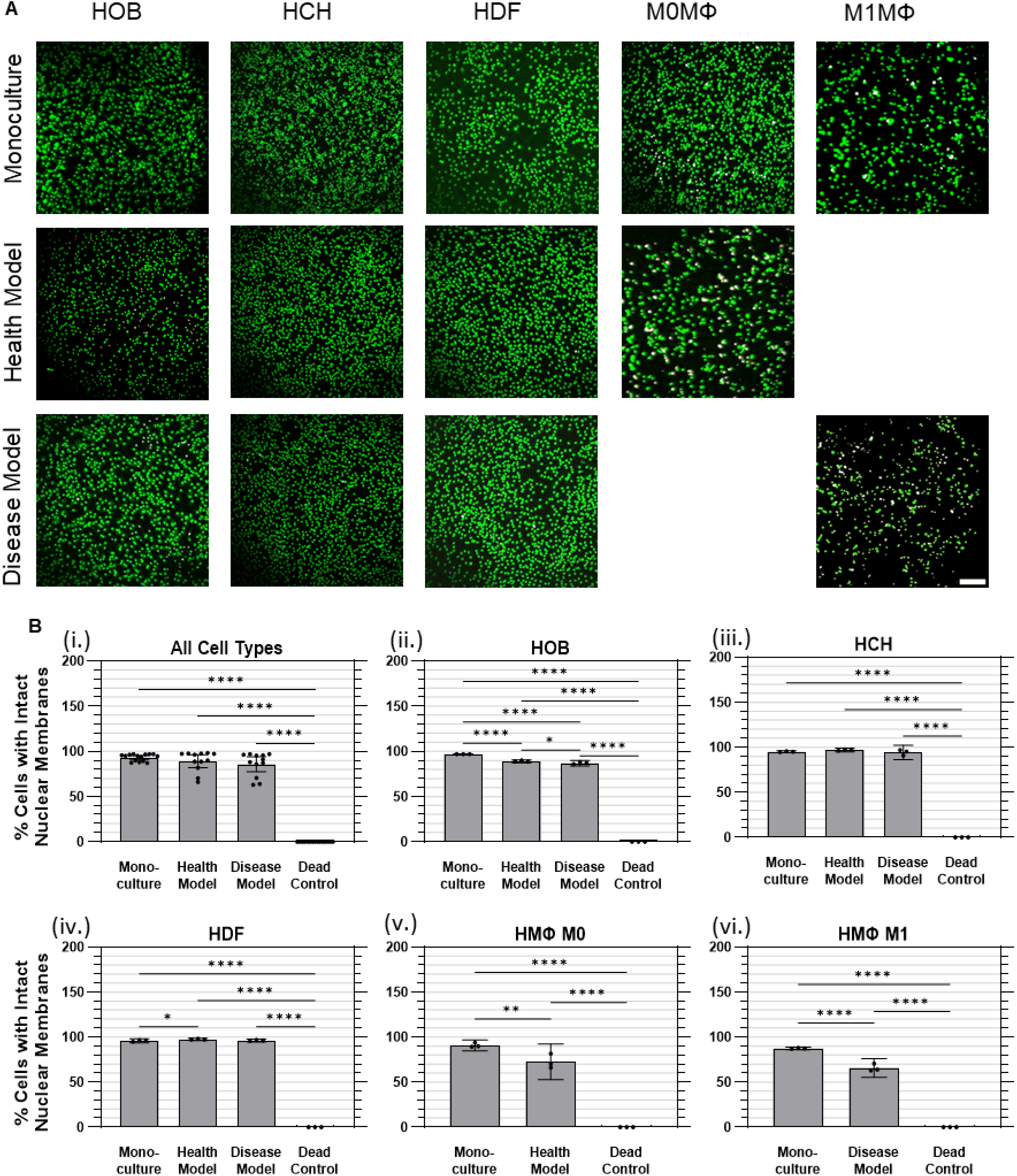
Cell viability was largely maintained at 24 hrs of co-culture in health and disease models. **(A)** Representative images of live (green) and dead (white) cell nuclei. **(B)** Quantified cell viability data for **(i)** Aggregated data across all cell types in monoculture, health co-culture, disease co-culture models, and dead controls; **(ii)** osteoblasts (HOB); **(iii)** chondrocytes (HCH); **(iv)** fibroblasts (HDF); **(v)** quiescent (M0) macrophages, i.e., Health Model; and **(vi)** pro-inflammatory (M1) macrophages, i.e., Disease Model. Data are mean ± 95% CI. ^*^*p* < 0.05, ^**^*p* < 0.01, ^****^*p* < 0.0001. Scale bar: 200 µm.

### Metabolic Activity Assessment

Metabolic activity, a reliable indicator of cell viability, was assessed using PrestoBlue™ HS Cell Viability Reagent (Invitrogen). (Boncler et al., 2014; Xu et al., 2015) This reagent, which relies on the reduction of resazurin to resorufin by metabolically active cells, provided a measure of cellular health. (Martín-Navarro et al., 2014) After a 2-hour incubation at 37°C, the resazurin was reduced by metabolically active cells, leading to a detectable increase in fluorescence, which was measured at an excitation/emission wavelength of 560/590 nm using a SpectraMax i3 plate reader (Molecular Devices, LLC., San Jose, CA)

### Cytotoxicity Assessment

To evaluate the cytotoxicity of co-culture conditions, cell membrane integrity was measured using the CyQUANT™ LDH Cytotoxicity Assay (Invitrogen). This assay quantifies the release of the intracellular protein lactate dehydrogenase (LDH) from damaged cells. Culture medium was collected from each cell type, and CyQUANT™ reagents were added according to the manufacturer’s protocol, and fluorescence from the enzymatic reaction was measured at 560/590 nm using a SpectraMax i3 plate reader.

### Statistical Analysis

Data from three independent experiments were analyzed for all statistical analyses. Effects of culture conditions were determined using one-way ANOVA with Tukey multiple comparisons post-hoc test with a single pooled variance. Effects were confirmed using a significance level of α=0.05. Data are represented as mean ± 95% confidence interval (95% CI). All statistical analyses were performed using GraphPad Prism 10.3.1.

## Results

### Viability

Cell viability was assessed using a ReadyProbes™ Cell Viability Imaging Kit to distinguish cells with intact nuclear membranes (i.e., live cells) from cells with disrupted nuclear membranes (i.e., dead cells), as described above. Dead controls were generated by treating cells that had been plated under monoculture conditions with 10X lysis buffer for 45 minutes to achieve complete cell death. Overall, when considering all cell types together within each culture condition, no measurable (i.e., statistically significant) differences in cell viability were observed between the monoculture (mean 93.3% ± 3.8% SD) and co-culture conditions (89.2% ± 11% health model, 85.9 ± 13% disease model), as indicated in Figure 4B(i) However, measurable differences (p<0.0001) were found between the dead control group (no detectable nuclei remained under any condition) and the monoculture and co-culture models.

For HOBs, cell viability in monoculture (97.0% ± 0.1%) and in health (89.2% ± 0.6%) and disease (87.0% ± 1.2%) co-culture models were all substantially higher than the dead control group (0%). A marginal but measurable difference in cell viability between health and disease models was observed (Figure 4B(ii)) For HCHs, no measurable difference in cell viability was found between monoculture (95.1% ± 0.5%) and health (97.2% ± 0.8%) or disease (94.4% ± 3.2%) co-culture models. However, all conditions showed substantially higher viability than the dead control group (0%); Figure 4B(iii)) HDF cell viability was marginally but measurably higher in the health co-culture model (97.6% ± 0.6%) compared to monoculture (96.0% ± 0.9%), while no measurable difference was observed between health and disease co-culture models (96.3% ± 0.6%). All conditions had higher viability compared to the dead control group (0%); (Figure 4B(iv)) For M0 quiescent macrophages, viability was measurably higher in monoculture (90.9% ± 2.4%) compared to the health co-culture model (72.8% ± 8.0%) and both conditions had substantially higher viability than the dead control group (0%); (Figure 4B(v)) For M1 pro-inflammatory macrophages, viability in monoculture (87.6% ± 0.6%) and disease co-culture model (65.8% ± 4.1%) were both lower than for other cell types, but were still substantially higher than the dead control group (0%), with a measurable difference between monoculture and disease co-culture models (Figure 4B(vi)).

### Metabolic Activity

Cell metabolic activity was assessed using the PrestoBlue™ assay, with levels of resorufin fluorescence relative to monoculture conditions (mean 1.00 ± 0.19 RFU [SD]) serving as an indicator of cell viability. Overall, the health co-culture model exhibited the highest metabolic activity, with a mean relative fluorescence of 5.89 ± 3.17 RFU, followed by the disease co-culture model at 5.28 ± 3.42 RFU. In contrast, the dead control group showed minimal metabolic activity, with a maximum fluorescence of 0.10 ± 0.03 RFU (Figure 5(A)).

**Figure 5:**
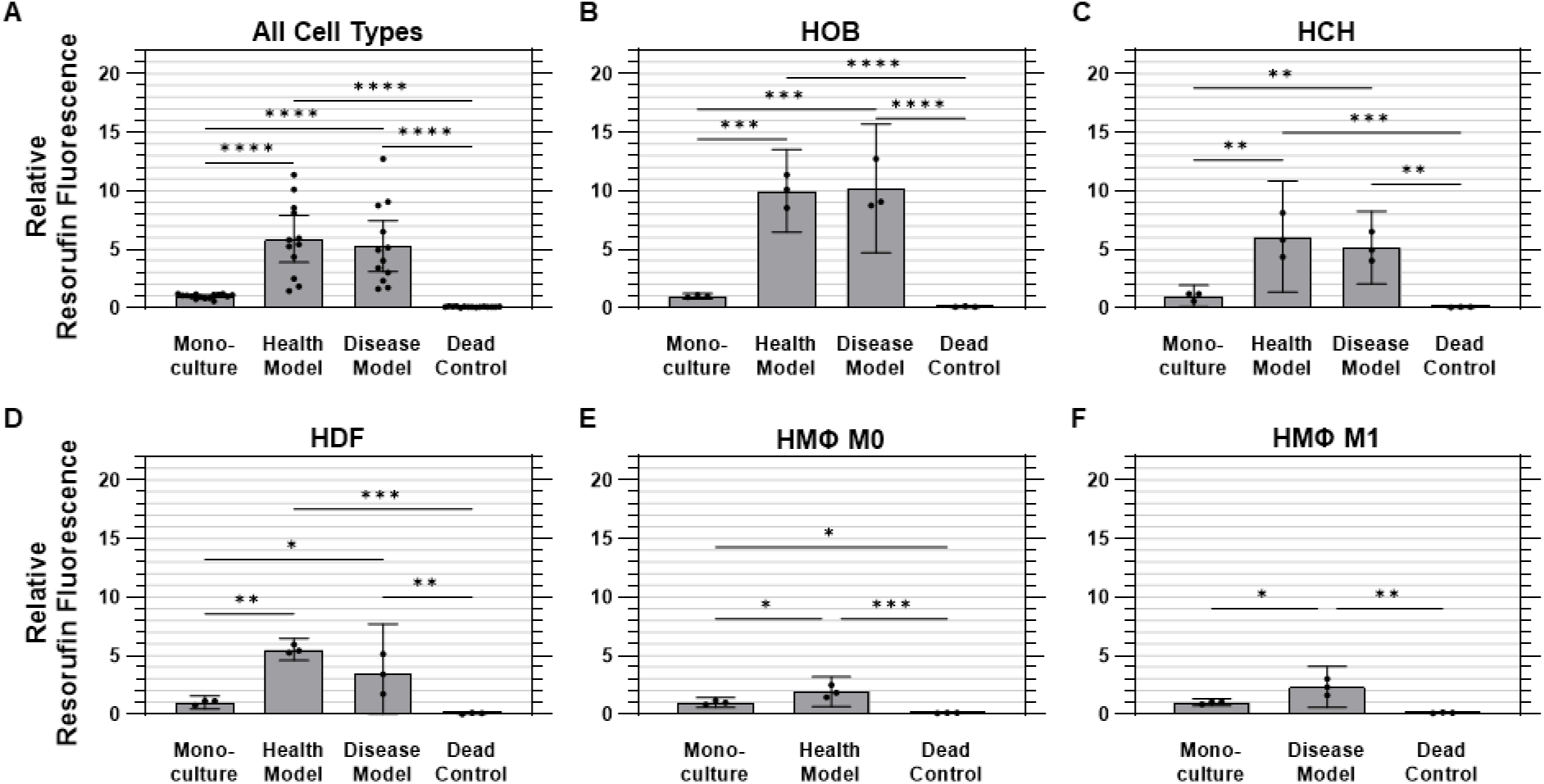
Metabolic activity was elevated under co-culture conditions. Quantified metabolic activity data (relative fluorescence) for **(A)** aggregated data across all cell types in monoculture, health co-culture, disease co-culture models, and dead controls; **(B)** osteoblasts (HOB); **(C)** chondrocytes (HCH); **(D)** fibroblasts (HDF); **(E)** quiescent (M0, i.e., Health Model) macrophages; and **(F)** pro-inflammatory (M1, i.e., Disease Model) macrophages. Data are mean ± 95% CI relative to monoculture. ^*^p<0.05, ^**^p<0.01, ^***^p<0.001, ^****^p<0.0001.

For HOBs, both co-culture models were found to have measurably higher metabolic activity than monoculture and dead control conditions. The health co-culture model showed a mean fluorescence of 10.0 ± 1.4 times those of monoculture HOBs (1.00 ± 0.09 RFU), while the disease co-culture model demonstrated a roughly equivalent value of 10.2 ± 2.2 times monoculture values. Dead controls showed minimal metabolic activity at 0.11 ± 0.03 RFU (Figure 5(B)) For HCHs, both co-culture models were also found to have measurably higher metabolic activity than monoculture (1.00 ± 0.4 RFU) and dead control (0.08 ± 0.03 RFU) conditions. The health co-culture model displayed the highest metabolic activity of all conditions, with a mean fluorescence of 6.09 ± 1.9 RFU, followed by the disease co-culture model at 5.16 ± 1.2 RFU, although the difference between these values was not found to be statistically measurable (Figure 5(C)) Both co-culture models were found to have measurably higher metabolic activity than monoculture (1.00 ± 0.2 RFU) and dead control (0.07 ± 0.04 RFU) conditions for HDFs as well. As was found for HOBs, the health co-culture model exhibited the highest metabolic activity for HDFs, with a mean fluorescence of 5.54 ± 0.4 RFU, followed by the disease co-culture model at 3.44 ± 1.7 RFU, with the difference between the two not found to be statistically measurable (Figure 5(D)) For M0 macrophages, the health co-culture model exhibited a mean metabolic activity of 1.92 ± 0.5 times that of monoculture (1.00 ± 0.2 RFU). The M0 dead controls exhibited minimal activity at 0.10 ± 0.02 RFU. All M0 conditions were found to be measurably different from one another (Figure 5(E)) M1 macrophages in the disease co-culture model displayed a mean metabolic activity (2.33 ± 0.7 RFU) measurably higher than those of M1 macrophages in either monoculture (1.00 ± 0.1 RFU) or dead control (0.13 ± 0.03 RFU) conditions (Figure 5(F)).

### Cytotoxicity

Cytotoxicity was evaluated using the lactate dehydrogenase (LDH) assay, which measures the release of intracellular LDH from cells as an indicator of compromised membrane integrity due to cell damage or death. No measurable differences were observed between monoculture and either co-culture model for any cell type, while all three culture conditions were found to release measurably less LDH into conditioned media than the cells in the dead controls for all cell types except HDFs, which were found to have an abnormally high degree of variability in the dead control samples. On the whole, cells cultured under monoculture conditions released 86.2% ± 8.3% less (i.e., 0.138 ± 0.083 times as much) LDH than dead control cells (1.00 ± 0.17) on average, while those in the health and disease co-culture models released 83.7% ± 10.9% and 87.4% ± 9.6% less LDH than dead control cells, respectively (Figure 6(A)).

**Figure 6:**
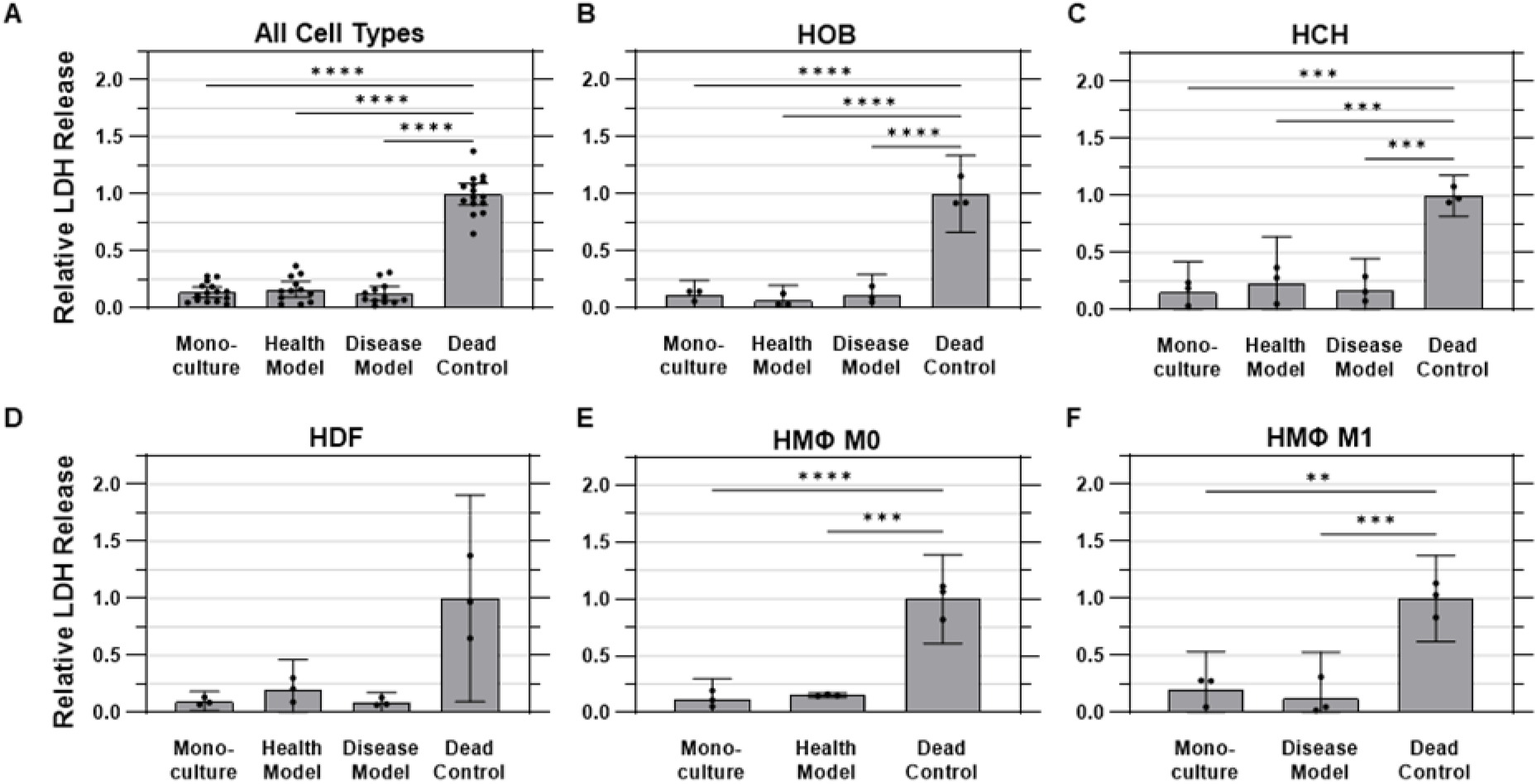
Membrane integrity was not substantially impacted by co-culture. Quantified LDH release into conditioned media, representing cell membrane integrity across various experimental conditions. **(A)** Aggregated data across all cell types in monoculture, health co-culture, disease co-culture models, and dead controls; **(B)** osteoblasts (HOB); **(C)** chondrocytes (HCH); **(D)** fibroblasts (HDF); **(E)** quiescent (M0, *i*.*e*., Health Model) macrophages; and **(F)** pro-inflammatory (M1, *i*.*e*., Disease Model) macrophages. Data are mean ± 95% CI relative to dead controls.

For HOBs, the dead control cells exhibited the highest LDH release levels, as expected, with a standard deviation of 10% of the mean value. The monoculture condition showed a substantially lower amount of LDH release, at 12% ± 5% as much as the dead control, while the health co-culture model had an even lower, but not statistically different, value of 6.5% ± 5.4% as much as the dead control. The disease model exhibited an LDH release of 11.2% ± 7.3% of dead control levels (Figure 6(B)) For HCHs, the dead control again had the highest normalized LDH release, with a standard deviation of 7% of the mean value. The health co-culture model exhibited LDH release of 23.4% ± 0.16% as much as dead control cells, followed by the disease co-culture model at 17.5% ± 11% and monoculture at 15.3% ± 11% of dead control values (Figure 6(C)) The dead control also displayed the highest amount of LDH release for HDFs, with a standard deviation of 36% of the mean value. The monoculture, health model, and disease model conditions exhibited LDH release levels of 9.8% ± 3%, 19.9% ± 11%, and 9.0% ± 3% as much as the dead control, respectively (Figure 6(D)) For quiescent (M0) macrophages, the dead control cells released the highest amount of LDH, with a standard deviation of 16% of the mean value. The monoculture and health co-culture models showed lower LDH release levels of 12.0% ± 7% and 15.4% ± 1% of the levels of the dead control, respectively (Figure 6(E)) For pro-inflammatory (M1) macrophages, the dead control once again exhibited the highest LDH release, with a standard deviation of 15% of the mean value. The monoculture and disease model conditions were observed to release 20.1% ± 13% and 12.7% ± 16% as much LDH as the dead control, respectively (Figure 6(F)).

## Discussion

Traditional *in vitro* models lack the complexity required to accurately recapitulate the multi-tissue nature of joint tissue interactions, thus limiting their relevance for OA research. While co-culture systems involving two joint cell or tissue types (e.g., bone/cartilage) have been explored, they may still fail to capture the broader intercellular crosstalk seen *in vivo*. Here, we address this limitation by introducing a co-culture model with osteoblasts, chondrocytes, fibroblasts, and quiescent (M0) or pro-inflammatory (M1) macrophages, thereby providing a more representative platform for studying the pathophysiology of OA.

The primary goal of this study was to develop a microfluidic co-culture model that integrates osteoblasts, chondrocytes, fibroblasts, and macrophages, mimicking the complex environment of the human joint. Our aim was to demonstrate that, by sharing paracrine signals ostensibly representative of the *in vivo* condition, cells can maintain a healthy condition similar to monoculture even when all cell types share one combined culture medium rather in lieu of their typical individualized medium. By providing a controlled co-culture environment, this study aims to establish a new foundation for studying osteoarthritis pathophysiology and evaluating potential therapeutic interventions.

We hypothesized that this system would maintain cell viability, support metabolic activity, and preserve membrane integrity for all cell types under both healthy and disease-mimicking conditions. Specifically, we predicted that viability would remain above 80% across all cell types, metabolic activity in co-culture would be comparable to monoculture conditions, and cytotoxicity would remain low and comparable to monoculture levels. Viability was successfully maintained across all conditions, with monoculture cells showing a mean viability of 93.3% ± 3.8%. The co-culture models exhibited slightly lower but comparable viability, with 89.2% ± 11% in the healthy model and 85.9% ± 13% in the disease model, all of which surpassed our hypothesized threshold of 80% viability (Figure 4B). Metabolic activity also met and exceeded expectations. Relative to monoculture (1.00 ± 0.19 RFU), metabolic activity was significantly elevated in the healthy co-culture model (5.89 ± 3.17 RFU) and disease co-culture model (5.28 ± 3.42 RFU), representing substantial increases in cellular metabolic function (Figure 5). Cytotoxicity, assessed through LDH release, showed no evidence of adverse effects due to co-culture conditions. Monoculture conditions released 86.2% ± 8.3% less LDH than dead control cells (1.00 ± 0.17 RFU), while the health and disease co-culture models released 83.7% ± 10.9% and 87.4% ± 9.6% less LDH, respectively (Figure 6).

While other models have co-cultured limited subsets of joint cells or tissues, such as osteoblasts with chondrocytes or cartilage with synovium, these models may fail to fully recapitulate the intercellular interactions present in the native joint microenvironment. To the best of our knowledge, no prior study has integrated four joint cell types into a single, shared microfluidic co-culture system. Furthermore, many of the previous studies relied on isolated tissue compartments or induced pluripotent stem cell (iPSC)-derived cells, which may not fully capture the physiological complexity of mature human cells. In contrast, our system incorporates primary human cells from bone, cartilage, synovium, and immune components, all of which interact within a shared medium under controlled flow conditions.

Lin et al. developed a multichamber bioreactor to model the bone-cartilage interface using human bone marrow stem cell (hBMSC)-derived constructs. Separate medium streams created distinct chondral and osseous microenvironments, enabling targeted chondrogenic and osteogenic differentiation over a four-week period. (Lin et al., 2014) In a subsequent study, they utilized a dual-flow bioreactor to replicate the bone-cartilage interface using induced pluripotent stem cell-derived mesenchymal progenitor cells (iMPCs). The iMPCs were encapsulated within photo-crosslinked gelatin scaffolds and exposed to distinct chondro- and osteo-inductive cytokine conditions, with chondrogenic medium flowing from the top and osteogenic medium from the bottom. (Lin et al., 2019) While both studies demonstrate progress in joint micro-physiological system development, the use of cells derived from a single source *in vitro* limit their biological relevance. Notably, neither study assessed cell viability or cytotoxicity under co-culture conditions, highlighting a key limitation that our model addresses by integrating four primary cell types from different donors with validated viability and cytotoxicity.

Mondadori et al. developed a microfluidic chip to model the synovium-cartilage interface, incorporating synovial and chondral compartments, an endothelial monolayer channel, and a dedicated synovial fluid channel. The system was populated with osteoarthritic synovial fibroblasts (SFbs) and articular chondrocytes (ACs) embedded in fibrin gel, while a perfusable endothelialized channel was established using human umbilical vein endothelial cells (HUVECs). This endothelial layer was designed to model vascularization and facilitate monocyte extravasation, simulating immune cell infiltration into the joint, a key feature in osteoarthritis pathology. To validate this, monocyte extravasation was assessed by perfusing TNF-α and chemokine gradients across the chip, which promoted migration through the endothelial layer into the synovial compartment. Further, OA synovial fluid from patient donors was used to trigger monocyte recruitment, highlighting the model’s relevance in studying immune responses under inflammatory conditions (Mondadori et al., 2021). While this system provides a sophisticated platform for investigating immune-cell recruitment and synovial-chondral interactions, it focuses on cells from synovial and chondral tissues only, without inclusion of bone-derived cells or macrophages. Additionally, no direct validation of long-term cell viability or cytotoxicity was provided under co-culture conditions, limiting its applicability for comprehensive joint modeling.

Rothbauer et al. developed a joint-on-a-chip model to explore the reciprocal tissue crosstalk between synovial and chondral organoids. Their system featured two spatially separated hydrogel-based 3D organoid constructs, with synovial organoids composed of fibroblast-like synoviocytes (FLS) derived from rheumatoid arthritis (RA) patients undergoing synovectomy and chondral organoids formed using commercially available human primary chondrocytes. The co-culture was facilitated by soluble signaling between the two compartments, enabling tissue-level communication to study joint pathophysiology. (Mario Rothbauer et al., 2021) This model also demonstrates a sophisticated approach to modeling joint interactions; however, it is limited by a focus on synovial and chondral tissues exclusively, lacking the inclusion of bone-derived cells or macrophages. It is further limited by a reliance on a mixture of healthy and diseased cells within a single co-culture system, lacking clear ‘healthy’ or ‘diseased’ reference states between which to make comparisons of cell behavior. Additionally, no data were reported on cell viability, cytotoxicity, or functional stability of the co-culture system. These limitations highlight the distinct advantage of the model presented in this study, which incorporates four key joint cell types from different tissue sources, with validated viability and lack of cytotoxicity, and offers a more comprehensive platform for studying joint diseases.

Kubosch et al. investigated the paracrine interactions between human synovial mesenchymal stem cells (SMSCs) and chondrocytes using an *in vitro* Transwell® monolayer co-culture system. (Kubosch et al., 2016) This approach enabled the examination of cellular crosstalk while maintaining physical separation between the two cell types. The SMSCs were isolated from synovial tissue collected during knee arthrotomy and arthroscopy procedures (n = 4, male/female 2/2). The chondrocytes were obtained from femoral heads during hip arthroplasty procedures (n = 6, male/female 5/1). While this study highlighted important paracrine interactions, it also lacked assessment of key functional metrics such as cell viability, cytotoxicity, or metabolic activity.

The present study represents a novel approach to OA modeling by co-culturing four distinct primary joint cell types—osteoblasts, chondrocytes, fibroblasts, and macrophages—within a unified microfluidic system with both healthy and diseased states governed by the cells within the system. This setup advances beyond prior models that typically incorporate only two cell types or rely on phenotypes derived from stem cells, offering a more physiologically relevant platform for joint research. The results presented here demonstrate the ability of the system presented in this study to sustain viability, metabolic activity, and membrane integrity across all cell types, providing a stable and functional co-culture environment. However, certain limitations should be noted. First, human dermal fibroblasts (HDFs) were used instead of synovial fibroblasts, which may not fully capture the functional role of synovial cells in an articular joint. Second, the co-culture period was limited to 24 hours, offering only a short-term perspective on cellular interactions, whereas longer-term studies may yield additional insights. Third, due to the technical requirements of the pre-conditioning period, primary chondrocytes were expanded and passaged prior to co-culture, which can alter their native phenotype. Despite these limitations, our model provides the foundation for a robust and versatile platform for OA research, supporting future studies on disease pathophysiology and therapeutic testing.

Future work will focus on expanding the capabilities of our microfluidic co-culture system. A key priority is the quantification of key biomarkers, such as matrix metalloproteinases (MMPs) and inflammatory cytokines, to characterize the healthy and diseased profiles of our co-culture design. This will enable a more precise characterization of the pathophysiological differences induced by co-culture conditions. Additional studies incorporating fibroblasts of synovial origin and phenotype-modulating biophysical cues to recapitulate native tissue architectures are expected to further extend the translational potential of this model in the future. Ultimately, we aim to test potential drug candidates targeting critical OA pathways, leveraging the system as a preclinical screening tool. These advancements will support the development of translationally viable OA therapies and further validate the system’s capacity as a predictive model for joint disease pathophysiology and therapeutic evaluation.

## Conclusion

In this study, we present a microfluidic co-culture system using ibidi µ-Slide I Luer microfluidic chips in which four cell types (osteoblasts, chondrocytes, fibroblasts, and two phenotypes of macrophages, M0 and M1) were integrated to simulate the complex cellular interactions of the human joint. Quiescent M0 macrophages represented the healthy model, while pro-inflammatory M1 macrophages simulated an inflammatory disease condition. This model will enable studies of different joint cells and the interactions between them in the context of both healthy and inflammatory environments. It is expected to provide a platform for OA and other rheumatic disease research, along with evaluating the potential efficacy of novel therapeutic approaches to treat these diseases. We observed that the present system-maintained cell viability, metabolic activity, and membrane integrity under co-culture conditions, revealed by NucBlue™/NucGreen™, PrestoBlue™, and LDH assays, respectively. No significant deleterious effects from crosstalk among cell types were observed, with increased metabolic activity consistently evident under co-culture conditions. These data could indicate that the model can mimic both healthy and diseased states without cytotoxicity and, therefore, is useful in studies of osteoarthritis pathophysiology. Furthermore, the stability of the system in inflammatory conditions indicated its potential for drug testing and therapeutic assessment of mainly treatments targeting inflammation. The model can be further optimized, but this microfluidic co-culture system paves the way toward more realistic models of the joint. Further studies could refine the system for high-throughput drug screening or investigate the use of this system for studying other inflammatory diseases or even other organs.

## Acknowledgements

This material is based upon work supported by the National Science Foundation under Grant No. 2517512.

## Disclaimer

SW is an inventor on a patent that pertains to the methods described in this publication for which he is entitled to receive royalties and/or equity. US Patent No. 12098354B2 was issued to the South Dakota Board of Regents. In addition, SW is a partner in a company, CellField Technologies, Inc., that has licensed related technology from the South Dakota Board of Regents.

## References Cited

Awad, H., Ajalik, R., Alenchery, R., Linares, I., Wright, T., Miller, B., & McGrath, J. (2023). Human tendon-on-a-chip for modeling vascular inflammatory fibrosis. Res Sq. 10.21203/rs.3.rs-3722255/v1

Banh, L., Cheung, K. K., Chan, M. W. Y., Young, E. W. K., & Viswanathan, S. (2022). Advances in organ-on-a-chip systems for modelling joint tissue and osteoarthritic diseases. Osteoarthritis Cartilage, 30(8), 1050–1061. 10.1016/j.joca.2022.03.012

Baxter, E. W., Graham, A. E., Re, N. A., Carr, I. M., Robinson, J. I., Mackie, S. L., & Morgan, A. W. (2020). Standardized protocols for differentiation of THP-1 cells to macrophages with distinct M(IFNγ+LPS), M(IL-4) and M(IL-10) phenotypes. J Immunol Methods, 478, 112721. 10.1016/j.jim.2019.112721

Boncler, M., Różalski, M., Krajewska, U., Podsędek, A., & Watala, C. (2014). Comparison of PrestoBlue and MTT assays of cellular viability in the assessment of anti-proliferative effects of plant extracts on human endothelial cells. J Pharmacol Toxicol Methods, 69(1), 9–16. 10.1016/j.vascn.2013.09.003

Cope, P. J., Ourradi, K., Li, Y., & Sharif, M. (2019). Models of osteoarthritis: the good, the bad and the promising. Osteoarthritis Cartilage, 27(2), 230–239. 10.1016/j.joca.2018.09.016

Domínguez-Oliva, A., Hernández-Ávalos, I., Martínez-Burnes, J., Olmos-Hernández, A., Verduzco-Mendoza, A., & Mota-Rojas, D. (2023). The Importance of Animal Models in Biomedical Research: Current Insights and Applications. Animals, 13(7), 1223. https://www.mdpi.com/2076-2615/13/7/1223

Dou, H., Wang, S., Hu, J., Song, J., Zhang, C., Wang, J., & Xiao, L. (2023). Osteoarthritis models: From animals to tissue engineering. J Tissue Eng, 14, 20417314231172584. 10.1177/20417314231172584

Goers, L., Freemont, P., & Polizzi, K. M. (2014). Co-culture systems and technologies: taking synthetic biology to the next level. J R Soc Interface, 11(96). 10.1098/rsif.2014.0065

Grässel, S., & Muschter, D. (2020). Recent advances in the treatment of osteoarthritis. F1000Res, 9. 10.12688/f1000research.22115.1

Guo, J., Huang, X., Dou, L., Yan, M., Shen, T., Tang, W., & Li, J. (2022). Aging and aging-related diseases: from molecular mechanisms to interventions and treatments. Signal Transduction and Targeted Therapy, 7(1), 391. 10.1038/s41392-022-01251-0

Haltmayer, E., Ribitsch, I., Gabner, S., Rosser, J., Gueltekin, S., Peham, J., Giese, U., Dolezal, M., Egerbacher, M., & Jenner, F. (2019). Co-culture of osteochondral explants and synovial membrane as in vitro model for osteoarthritis. PLOS ONE, 14(4), e0214709. 10.1371/journal.pone.0214709

He, Y., Li, Z., Alexander, P. G., Ocasio-Nieves, B. D., Yocum, L., Lin, H., & Tuan, R. S. (2020). Pathogenesis of Osteoarthritis: Risk Factors, Regulatory Pathways in Chondrocytes, and Experimental Models. Biology (Basel), 9(8). 10.3390/biology9080194

Hofer, M., & Lutolf, M. P. (2021). Engineering organoids. Nat Rev Mater, 6(5), 402–420. 10.1038/s41578-021-00279-y

Kao, C.-Y., Mills, J. A., Burke, C. J., Morse, B., & Marques, B. F. (2023). Role of Cytokines and Growth Factors in the Manufacturing of iPSC-Derived Allogeneic Cell Therapy Products. Biology, 12(5), 677. https://www.mdpi.com/2079-7737/12/5/677

Kubosch, E. J., Heidt, E., Bernstein, A., Böttiger, K., & Schmal, H. (2016). The trans-well coculture of human synovial mesenchymal stem cells with chondrocytes leads to self-organization, chondrogenic differentiation, and secretion of TGFβ. Stem Cell Research & Therapy, 7(1), 64. 10.1186/s13287-016-0322-3

Li, Z. A., Sant, S., Cho, S. K., Goodman, S. B., Bunnell, B. A., Tuan, R. S., Gold, M. S., & Lin, H. (2023). Synovial joint-on-a-chip for modeling arthritis: progress, pitfalls, and potential. Trends Biotechnol, 41(4), 511–527. 10.1016/j.tibtech.2022.07.011

Lin, H., Lozito, T. P., Alexander, P. G., Gottardi, R., & Tuan, R. S. (2014). Stem cell-based microphysiological osteochondral system to model tissue response to interleukin-1β. Mol Pharm, 11(7), 2203–2212. 10.1021/mp500136b

Lin, Z., Li, Z., Li, E. N., Li, X., Del Duke, C. J., Shen, H., Hao, T., O’Donnell, B., Bunnell, B. A., Goodman, S. B., Alexander, P. G., Tuan, R. S., & Lin, H. (2019). Osteochondral Tissue Chip Derived From iPSCs: Modeling OA Pathologies and Testing Drugs. Front Bioeng Biotechnol, 7, 411. 10.3389/fbioe.2019.00411

Loeser, R. F., Goldring, S. R., Scanzello, C. R., & Goldring, M. B. (2012). Osteoarthritis: a disease of the joint as an organ. Arthritis Rheum, 64(6), 1697–1707. 10.1002/art.34453

Long, H., Liu, Q., Yin, H., Wang, K., Diao, N., Zhang, Y., Lin, J., & Guo, A. (2022). Prevalence Trends of Site-Specific Osteoarthritis From 1990 to 2019: Findings From the Global Burden of Disease Study 2019. Arthritis Rheumatol, 74(7), 1172–1183. 10.1002/art.42089

Makarcyzk, M. J., Li, Z. A., Yu, I., Yagi, H., Zhang, X., Yocum, L., Li, E., Fritch, M. R., Gao, Q., Bunnell, B. A., Goodman, S. B., Tuan, R. S., Alexander, P. G., & Lin, H. (2023). Creation of a Knee Joint-on-a-Chip for Modeling Joint Diseases and Testing Drugs. J Vis Exp(191). 10.3791/64186

Malfait, A. M., & Little, C. B. (2015). On the predictive utility of animal models of osteoarthritis. Arthritis Res Ther, 17(1), 225. 10.1186/s13075-015-0747-6

Martín-Navarro, C. M., López-Arencibia, A., Sifaoui, I., Reyes-Batlle, M., Cabello-Vílchez, A. M., Maciver, S., Valladares, B., Piñero, J. E., & Lorenzo-Morales, J. (2014). PrestoBlue® and AlamarBlue® are equally useful as agents to determine the viability of Acanthamoeba trophozoites. Exp Parasitol, 145 Suppl, S69-72. 10.1016/j.exppara.2014.03.024

McNerney, M. P., Doiron, K. E., Ng, T. L., Chang, T. Z., & Silver, P. A. (2021). Theranostic cells: emerging clinical applications of synthetic biology. Nature Reviews Genetics, 22(11), 730–746. 10.1038/s41576-021-00383-3

Mondadori, C., Palombella, S., Salehi, S., Talò, G., Visone, R., Rasponi, M., Redaelli, A., Sansone, V., Moretti, M., & Lopa, S. (2021). Recapitulating monocyte extravasation to the synovium in an organotypic microfluidic model of the articular joint. Biofabrication, 13(4). 10.1088/1758-5090/ac0c5e

Palasantzas, V. E. J. M., Tamargo-Rubio, I., Le, K., Slager, J., Wijmenga, C., Jonkers, I. H., Kumar, V., Fu, J., & Withoff, S. (2023). iPSC-derived organ-on-a-chip models for personalized human genetics and pharmacogenomics studies. Trends in Genetics, 39(4), 268–284. 10.1016/j.tig.2023.01.002

Perisin, M. A., & Sund, C. J. (2018). Human gut microbe co-cultures have greater potential than monocultures for food waste remediation to commodity chemicals. Scientific Reports, 8(1), 15594. 10.1038/s41598-018-33733-z

Pirosa, A., Tankus, E. B., Mainardi, A., Occhetta, P., Dönges, L., Baum, C., Rasponi, M., Martin, I., & Barbero, A. (2021). Modeling In Vitro Osteoarthritis Phenotypes in a Vascularized Bone Model Based on a Bone-Marrow Derived Mesenchymal Cell Line and Endothelial Cells. Int J Mol Sci, 22(17). 10.3390/ijms22179581

Rothbauer, M., Byrne, R. A., Schobesberger, S., Olmos Calvo, I., Fischer, A., Reihs, E. I., Spitz, S., Bachmann, B., Sevelda, F., Holinka, J., Holnthoner, W., Redl, H., Toegel, S., Windhager, R., Kiener, H. P., & Ertl, P. (2021). Establishment of a human three-dimensional chip-based chondro-synovial coculture joint model for reciprocal cross talk studies in arthritis research. Lab Chip, 21(21), 4128–4143. 10.1039/d1lc00130b

Rothbauer, M., Byrne, R. A., Schobesberger, S., Olmos Calvo, I., Fischer, A., Reihs, E. I., Spitz, S., Bachmann, B., Sevelda, F., Holinka, J., Holnthoner, W., Redl, H., Toegel, S., Windhager, R., Kiener, H. P., & Ertl, P. (2021). Establishment of a human three-dimensional chip-based chondro-synovial coculture joint model for reciprocal cross talk studies in arthritis research [10.1039/D1LC00130B]. Lab on a Chip, 21(21), 4128–4143. 10.1039/D1LC00130B

Salgado, C., Jordan, O., & Allémann, E. (2021). Osteoarthritis In Vitro Models: Applications and Implications in Development of Intra-Articular Drug Delivery Systems. Pharmaceutics, 13(1). 10.3390/pharmaceutics13010060

Swearengen, J. R. (2018). Choosing the right animal model for infectious disease research. Animal Model Exp Med, 1(2), 100–108. 10.1002/ame2.12020

Tong, L., Yu, H., Huang, X., Shen, J., Xiao, G., Chen, L., Wang, H., Xing, L., & Chen, D. (2022). Current understanding of osteoarthritis pathogenesis and relevant new approaches. Bone Research, 10(1), 60. 10.1038/s41413-022-00226-9

Weiskirchen, S., Schröder, S. K., Buhl, E. M., & Weiskirchen, R. (2023). A Beginner’s Guide to Cell Culture: Practical Advice for Preventing Needless Problems. Cells, 12(5). 10.3390/cells12050682

Xie, X., Zhu, J., Hu, X., Dai, L., Fu, X., Zhang, J., Duan, X., & Ao, Y. (2018). A co-culture system of rat synovial stem cells and meniscus cells promotes cell proliferation and differentiation as compared to mono-culture. Scientific Reports, 8(1), 7693. 10.1038/s41598-018-25709-w

Xu, M., McCanna, D. J., & Sivak, J. G. (2015). Use of the viability reagent PrestoBlue in comparison with alamarBlue and MTT to assess the viability of human corneal epithelial cells. J Pharmacol Toxicol Methods, 71, 1–7. 10.1016/j.vascn.2014.11.003

Zaki, S., Blaker, C. L., & Little, C. B. (2022). OA foundations – experimental models of osteoarthritis. Osteoarthritis and Cartilage, 30(3), 357–380. 10.1016/j.joca.2021.03.024

Zhou, T., Yuan, Z., Weng, J., Pei, D., Du, X., He, C., & Lai, P. (2021). Challenges and advances in clinical applications of mesenchymal stromal cells. Journal of Hematology & Oncology, 14(1), 24. 10.1186/s13045-021-01037-x

